# Gyne production is regulated by the brood in a social bee (*Bombus impatiens*)

**DOI:** 10.1101/2022.07.04.498320

**Authors:** Etya Amsalem, Priscila K. F. Santos, Ella Messner, Cameron S. Murray

## Abstract

Sexual production in social insects marks the peak of colony development, yet the mechanisms regulating it remain unclear. We investigated the role of brood in colony development, worker reproduction, and sexual production in *Bombus impatiens*. While larvae are known to reduce worker egg-laying and enhance the queen’s reproductive inhibition, these effects were previously tested only in small groups. We manipulated brood size in full-sized, young colonies by doubling or removing brood and monitored development. Colonies with doubled brood produced significantly more gynes, independent of the number of workers, while reduced-brood colonies exhibited a non-significant increase in male production that was driven by colony size. Worker ovary activation was lower in double-brood colonies, with no change in aggression. A follow-up experiment directly testing the effect of colony size showed that higher worker density led to higher ovary activation in workers but did not affect sexual production. These results suggest that brood strongly influences colony development and sexual production, possibly reflecting an extended phenotype of the queen, whereas worker ovary activation appears to be a more flexible process influenced by either brood presence or colony size. Understanding brood dynamics may be key to understanding the evolution of female castes in social insects.

## Introduction

Social insects can form giant colonies; however, most individuals are sterile helpers produced to support the generation of a limited number of sexuals. Annual social species like bumble bees (1) and social wasps (2) slowly build up the worker force to support sexual production in a single event via a strategy called “bang-bang" theory (an abrupt shift from producing workers to sexuals) (3), whereas perennial species like honey bees and many species of ants alternate between cycles of worker and sexual production. In hymenopteran social insects, gynes (new queens) are morphologically, physiologically, and behaviourally different from female workers (4) and their life trajectory is often determined during larval development (5, 6). The determination of female caste early in development sets the foundation for the reproductive division of labour among females, and therefore, for the social organization (6). Timing the production of sexuals, and particularly of gynes, is critical for synchronizing sexual emergence with mating and floral availability and is regulated carefully to prevent loss of fitness. However, the factors triggering this event remain mostly unknown (7), and studying them is of significant importance to understanding social behaviour, resource allocation, and the evolution of female castes.

Research about the transition to sexual production in social insects has focused on a limited number of species, mostly bumble bees, honeybees, social wasps, and some ants. It is best understood through the lens of optimal resource allocation theory (8, 9), which predicts that colonies should initiate investment in sexuals once sufficient resources and workforce are secured to support reproductive success. Empirical studies support this framework but highlight the dominant role of internal sociobiological cues over environmental factors (7, 10). For instance, mathematical models and experimental data in bumble bees suggest that colony size and worker accumulation influence the optimal timing of reproduction (8, 11), yet actual triggers are often mediated by queen signalling. Queen presence (12) and age (13), pheromonal cues, and the type of brood in the colony (14, 15) regulate the onset of gyne production in *Bombus terrestris*. In ants such as *Linepithema humile* and *Aphaenogaster senilis*, queen control and pheromones, colony size and environmental seasonality jointly modulate sexual production (16–20), while in other species it might be seasonal (*Apis mellifera)* (21) or even random (*Formica exsecta*) (10). Across taxa, the lack of influence from worker age, mortality, or density (13, 22, 23) emphasize that sexual production is less a product of environmental feedback and more a colony-driven process shaped by the need to maximize inclusive fitness.

Recent work has begun to highlight the role of brood in shaping colony-level reproductive dynamics, particularly in Hymenoptera (24). Brood at various developmental stages, including larvae and pupae in *Apis mellifera* (25–27), larvae in *Novomessor cockerelli* (28) and *Ooceraea biroi* (29), eggs in *Camponotus floridanus* (30), and larvae and pupae in *Bombus impatiens* (31) has been shown to influence female reproduction and behaviour. Brood signals not only suppress worker reproduction but also regulate foraging behaviour, shifting workers from nursing to foraging roles and increasing foraging effort (27, 32–34). In several species of ants, different stages of brood were shown to stimulate egg production in fertile females. For example, late instar larvae in fire ants (35), and the presence of larvae in pharaoh’s ants drive queen reproduction (36). It was further suggested that the brood-to-worker ratio may influence gyne production in bumble bees (37), that workers can recognize and respond differently to sexual versus worker brood in Argentine ants (20), and that late-stage larvae in *Monomorium pharaonis* ants affect gyne production (38); however, the broader potential role of brood as a key trigger for sexual production has largely been overlooked.

Here we examined how the amount of brood affects colony development and demography using full size colonies of bumble bees (*Bombus impatiens)*. Bumble bees are an excellent system to examine these questions since they are annual and semelparous. The life cycle of the colony starts with a single, mated queen that lays eggs following a winter diapause. Initially, the queen performs all the tasks in the colony but she switches exclusively to egg laying once the first worker emerges (1). The queen monopolizes reproduction for approximately 4-5 weeks from the first worker emergence but loses dominance as the colony grows and transitions into the competition phase. During this phase, which highly correlates with the timing of gyne production (39), workers compete with the queen and among themselves over male production by exhibiting aggressive behaviour, oophagy and egg laying (40, 41). Gynes and males are typically produced towards the end of the season, though males can be produced earlier (42). Colonies differ substantially in the number and type of sexuals they produce, with some colonies specializing in producing gynes and others in producing males (43). This split sex ratio is partially explained by the diapause regime the queen experiences prior to funding a colony (43).

Recent studies in *Bombus impatiens* show that (a) young larvae decrease while pupae increase worker egg laying (31); (b) the impact of the queen on worker ovary activation is stronger when combined with the brood (44); and (c) the queen pheromonal secretion is only effective when combined with brood (45). Altogether, these studies demonstrate how significant is the brood to the social organization. To test the effects of brood on worker reproduction and sexual production, we manipulated brood amount in 16 queenright, young colonies prior to the onset of gyne production by transferring all brood from one colony to another. This resulted in five colonies with no brood, five colonies with double brood, and six unmanipulated colonies at the start of the experiment. We examined colony growth, aggressive behaviour toward and by the queen, worker ovary activation, and the production of egg batches, brood, workers, and sexuals over 26 days. From day 27 onward, we continued monitoring all adults emerging in the colonies until the emergence of the last bee. In a follow-up experiment, we monitored an additional 15 colonies, each restricted to either 30 or 90 workers, for 54 days using the same protocol. We hypothesized that an increased amount of brood would decrease worker ovarian activation and aggression and support the production of more gynes. We further hypothesized that, with the same amount of brood, colony size would affect worker ovary activation but not gyne production.

## Material and Methods

### Bumble bee rearing

A total of 31 *Bombus impatiens* colonies were used in this study. All colonies were obtained from Koppert Biological Systems (Howell, MI, USA) and arrived a few weeks after the emergence of the first worker, containing a queen, several dozen workers, and brood at various stages. Colonies were maintained in the laboratory under constant darkness, 60% relative humidity, and a temperature of 28–30 °C. They were provided with unlimited 60% sucrose solution and fresh pollen collected by honeybees, purchased from Koppert. All handling was performed under red light.

### Experimental design

In the first experiment (Figures 1-4, S1-2), 16 colonies of approximately equal wet mass and similar numbers of workers and brood were assigned to three treatments: in five colonies, all brood was transferred to another five colonies, creating ’no brood’ and ’double brood’ groups, respectively. The remaining six colonies were left unmanipulated and served as controls. Because precise brood counts were not possible, ‘double’ refers to receiving brood from two colonies. All colonies retained their original queen and workers. The experiment was conducted in two consecutive replicates (‘rounds’), each including eight colonies (three double brood, three no brood, and two controls in the first round; the remainder in the second). For 26 days post-manipulation, we monitored colony growth and mass (see below), observed aggression involving the queen, sampled workers for ovary activation at five timepoints, and recorded the number of new egg batches daily. On day 26, we removed all bees and brood, excluding pupae. We then counted the numbers of eggs, larvae, and pupae, and measured larval body mass. Pupae were left in the colonies until adult emergence, at which point emerging individuals were sexed and counted and added to the total number of individuals in each group.

**Figure 1:**
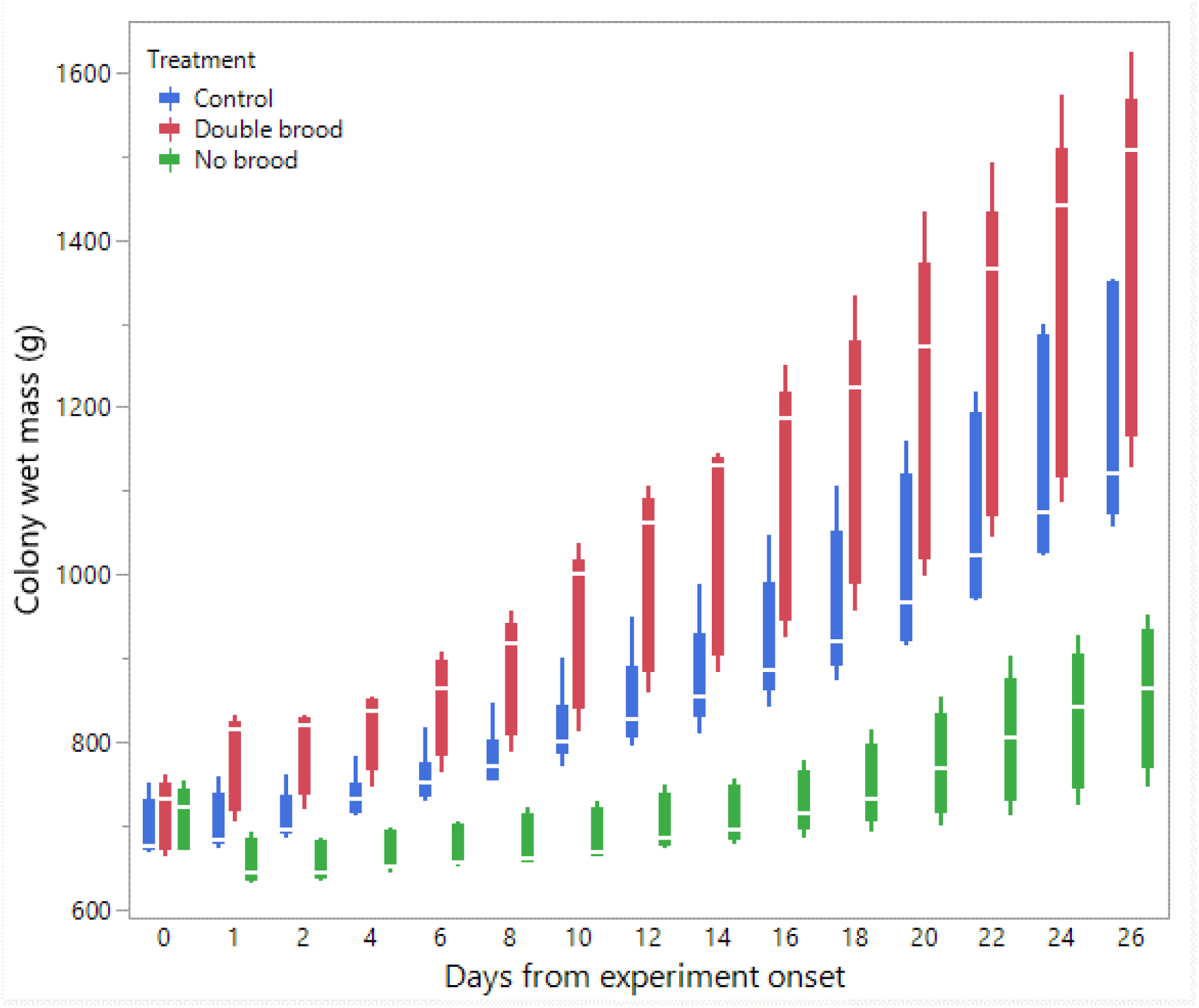
The average colony wet mass throughout the experiment in the three treatment groups: no brood, double brood, and control. Timepoints 0 and 1 refer to the colony wet mass before and after brood manipulation, respectively.

In the second experiment (Figures 5-8, S3), we used 15 colonies. For each colony, all individuals were removed, the brood wet mass was measured, and then the queen was returned along with either 30 or 90 workers, resulting in two treatments: 30 workers (n = 8) and 90 workers (n = 7). For 54 days following the manipulation, we repeated the same measurements as in the first experiment, excluding observations of aggressive behaviour. Previous studies have shown that even small colonies with as few as 20 workers are capable of producing gynes (13).

### Control for colony growth

To account for differences in colony growth during the first experiment (i.e., colonies without brood could not immediately produce new workers), newly emerged workers (<24 hours old) were collected and redistributed daily. These workers were equally divided among all colonies to ensure comparable growth regardless of initial brood quantity. In each round, a minimum of 8 and a maximum of 67 workers were collected daily and evenly distributed across the colonies. When the number of workers did not divide evenly by eight, the remaining individuals were allocated in the following order: no brood, control, and then double brood colonies, with balancing adjustments made on the consecutive day. In the second experiment, colony size was maintained at 30 or 90 workers by removing workers daily.

Although these methods were generally effective, perfect standardization was not possible due to the large number of workers per colony. Therefore, we also examined and controlled for the effects of the initial, final, and total number of workers on the measured variables (see Results).

### Colony mass

In the first experiment, colonies were weighed twice at the start, before and after the social manipulation, and then every other day throughout the experiment using an electronic scale. In the second experiment, colonies were first emptied of adults and food to obtain the actual brood mass, then weighed with the adults, and subsequently weighed daily. Weighing was performed by placing the entire colony on a scale, a method that is non-intrusive and not stressful to the bees. The recorded mass includes the bees, brood, cells, and the plastic box housing the colony, but excludes the sugar reservoir. All colonies were kept in their original identical Koppert plastic boxes.

### Aggressive behaviour

Colonies of the first experiment were video recorded every other day to assess aggression towards and by the queen. Each colony was recorded for 20 minutes between 9 am and 1 pm. Approximately 70 hours of video footage were analysed by an observer blind to the experimental design and hypotheses. Three behaviours were identified and summed: (1) attack: overt aggression including pulling, climbing, dragging, or attempted stinging; (2) darting: a sudden movement toward another bee without physical contact; and (3) humming: a series of short wing vibrations directed at another bee while in motion (46–48). Behaviours performed by the queen toward workers (“queen aggression”) and by workers toward the queen (“worker aggression”) were analysed separately and reported as the average number of behaviours observed per 20-minute recording.

### Worker ovarian activation

In the first experiment, 10–15% of the workers in each colony were collected at five timepoints (days 1, 7, 13, 19, and 26), and a subset (5–10 workers per colony per timepoint) was dissected to assess ovary activation. The number of workers removed was negligible relative to the total colony size and did not meaningfully affect colony dynamics. Similar sampling approaches have been used in previous studies and are considered representative of the workers’ overall reproductive state (49–51). Since workers were not individually tagged and newly emerged callows were redistributed daily, some dissected individuals may have originated from other colonies. However, as these workers were added upon emergence, their reproductive behaviour is assumed to align with that of the host colony (48). In the second experiment, a subset of the workers removed daily was dissected at 9 timepoints (approximately every 5–10 days). To measure terminal oocyte size, the abdomen was dissected under a stereomicroscope, and both ovaries were placed in a drop of water. The three largest oocytes were measured using a stereomicroscope ruler, and their lengths were averaged (52). Results are presented in millimetres.

### Larval body mass

At the end of both experiments, larvae were weighed individually using an electronic scale. Mass distribution within each colony is presented for all larvae with a body mass greater than 0.1 g. This cutoff was selected because caste-related differences in body mass (queen vs. worker) become detectable only at the third instar, which corresponds to approximately 0.1 g (53). Although the precise mechanism of caste determination in *B. impatiens* is unknown, it is unlikely to occur earlier than in *B. terrestris* (54). Our goal was to examine whether larvae were distributed bimodally, reflecting the production of both workers/males and gynes, or unimodally, reflecting the production of workers/males only.

### Statistical analyses

Statistical analyses and visualization of the main text figures were performed using JMP 18 Pro. Colony wet mass (Figures 1 and 5) was examined using a linear mixed model (LMM) with repeated measures. The model included treatment (control, double brood, no brood), day and their interaction as fixed effects, ‘round’ as a random factor, with colony ID as the repeated subject and a AR(1) covariance structure to account for correlations across time. A Generalized Linear Mixed Model (GLMM) was conducted to examine the effects of treatment, day and their interaction on the number of egg batches laid per day in Figures 2 and 6 and the count of aggression in Figure S1. The model specified a negative binominal distribution for the response variable and included ‘round’ and colony ID as random effects. The effect of treatment, day and interaction on ovarian activation (Figures 3 and 7) was examined using LMM, with ‘round’ and ‘colony ID’ included as random factors. The number of eggs, larvae, pupae, males, gynes and workers at the end of the experiment, as well as the initial and total number of workers and the initial mass of brood (Figures 4 and 8) were tested using GLMM. We included ‘round’ and colony ID as random factors, and the initial and total number of workers as covariates, whenever possible (see results for details). We ran the models with the following distributions: normal for eggs, larvae and pupae, lognormal for the initial, end and total number of worker and negative binomial for gynes and males. The initial mass of brood was fitted with a normal distribution. Kaplan–Meier survival analysis was used to compare the timing of gyne production across treatments, with differences between groups assessed using the Log-Rank test. To test the effect of treatment on the bimodality of larval body mass distribution, we use the ACR method. Statistical significance was accepted at α=0.05. All tests were followed up by multiple comparisons (Tukey post hoc for all pairwise comparisons). Data are presented as box plots excluding Figures 2 and 7 which show individual data points along with a non-parametric regression fit.

**Figure 2:**
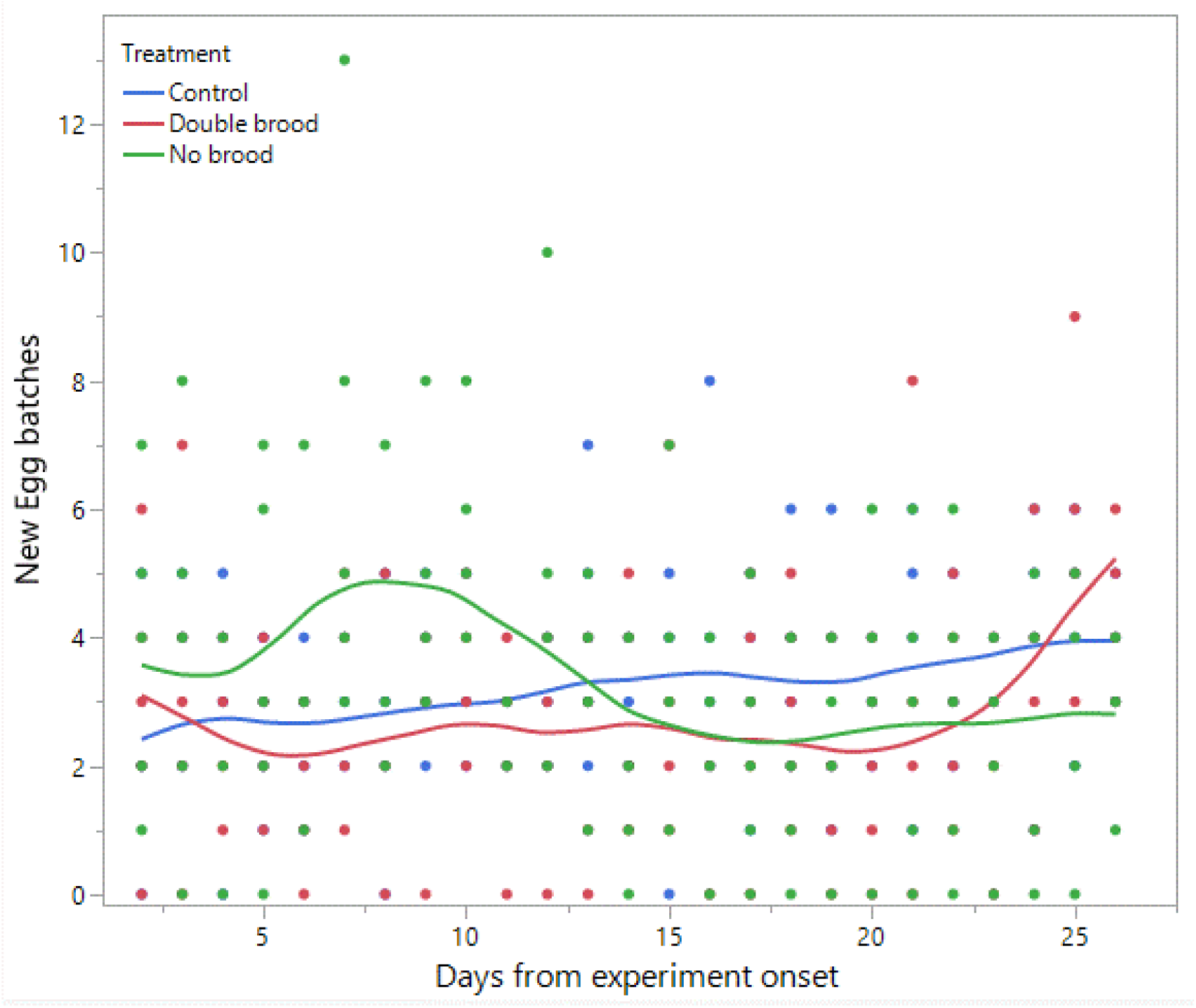
The number of egg batches laid per day throughout the experiment in the three treatment groups: no brood, double brood, and control.

**Figure 3:**
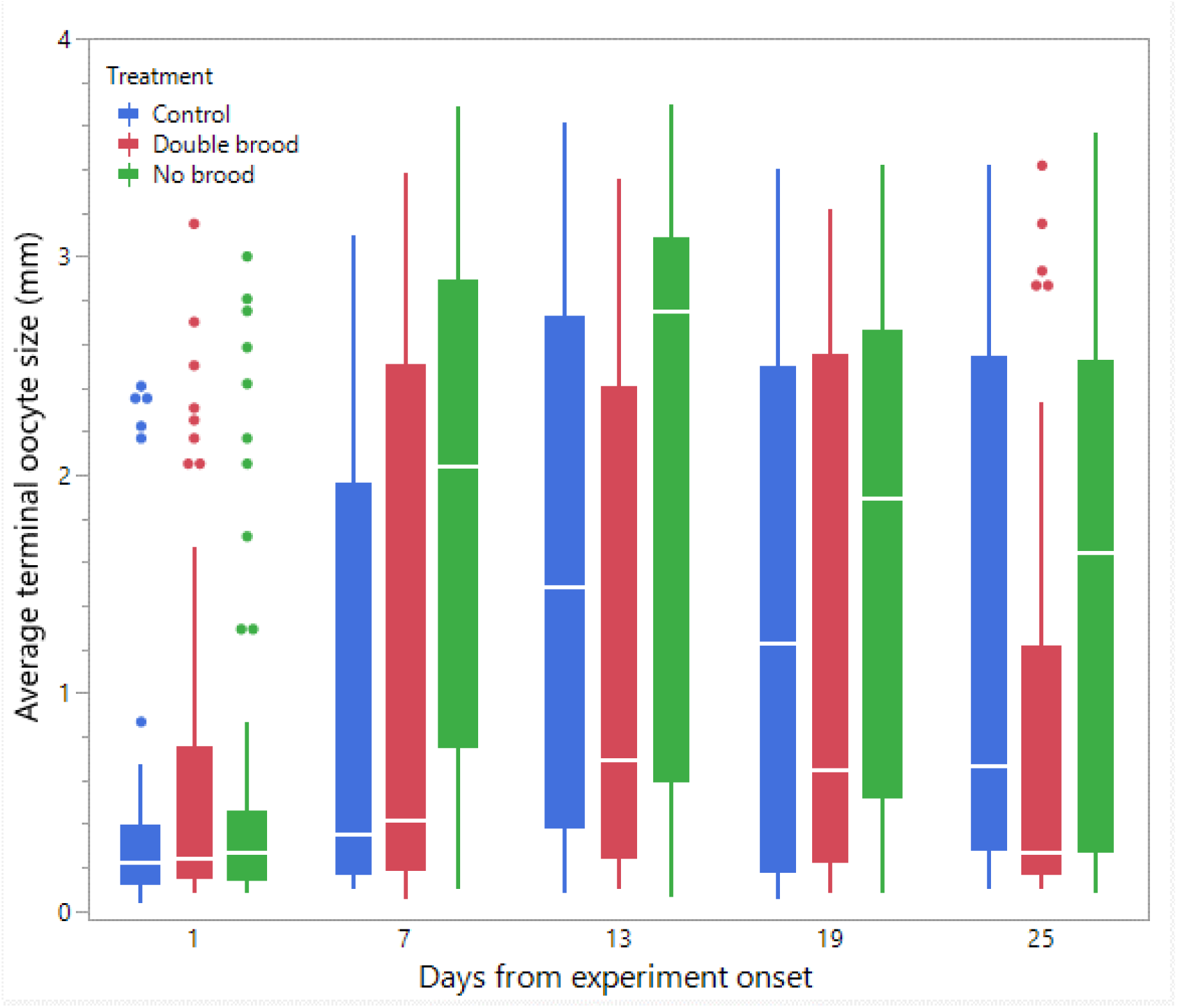
The average terminal oocyte size in workers in the three treatment groups: no brood, double brood, and control. Oocyte size was measured in five timepoints throughout the experiment using a subset of workers from each colony.

## Results

### Colony wet mass (Figure 1)

The analysis revealed a significant effect of treatment (F_2,12.9_=22.4, p<0.001 followed up by Tukey post-hoc test p<0.001 for no brood vs. the other treatments), day (F_1,148.9_=557.3, p<0.001) and the interaction between them (F_2,148.9_=67.6, p<0.001). All colonies gained mass over time, reflecting the increase in worker populations and brood. However, colonies in the double brood group exhibited significantly greater mass gain over time compared to the control group, while colonies in the no brood group gained significantly less mass over the same period.

### Production of new egg batches (Figure 2)

Neither treatment (F_2, 5.3_=0.34, p=0.32) nor day (F_1, 265.5_=0.98, p=0.32) had a significant main effect; however, their interaction was significant (F_2, 262.5_=12.6, p<0.001), indicating that colonies in the no brood treatment exhibited a modest increase in egg laying shortly after the onset of the experiment, peaking around day 7. This pattern likely reflects an immediate compensatory response to brood removal. Modest increases in egg laying were observed in the other two treatments at later time points, likely indicating the onset of worker reproduction.

### Workers’ ovary activation (Figure 3)

The model showed a significant main effect of treatment (F_2,12.2_= 4.26, p=0.039 followed by post hoc Tukey p=0.03 for no brood vs. double brood), day (F_4,731.2_= 22.79, p<0.0001 followed by Tukey test p<0.001 for d1 vs. all days and p=0.002 for d13 vs d25) and the interaction between them (F_8,731.3_= 2.6, p=0.008) indicating that treatment effects varied across time. The interaction estimates further highlighted no differences across treatments on day 1 and a pronounced increase in the no brood group by day 7. The no brood reached the highest observed value on day 13 (estimate = 2.16, 95% CI: 1.64 to 2.67), exceeding both control (1.62, 95% CI: 1.11 to 2.13) and double brood (1.17, 95% CI: 0.66 to 1.68). Although responses declined slightly on days 19 and 25, the no brood group consistently maintained higher values than the other treatments.

### The total number of brood and adults (Figure 4)

The number of workers at the end (Figure 4B) did not vary between treatments (F_2,1_=40.2, p=0.11), but both the initial number of workers (F_2,12.1_=5.66, p=0.01 followed by Tukey post-hoc test p=0.02 for double brood vs. control and p=0.04 for double brood vs. no brood, Figure 4A) and the total number of workers (F_2,9.4_=7.17, p=0.01 followed by Tukey post-hoc test p=0.01 for double brood vs. no brood, Figure 4C) were affected by the treatment. Therefore, to test the effect of treatment on the number of brood, males, and gynes, we included in the subsequent models both the random effects (round, colony ID) and the initial and total number of workers as covariates.

**Figure 4:**
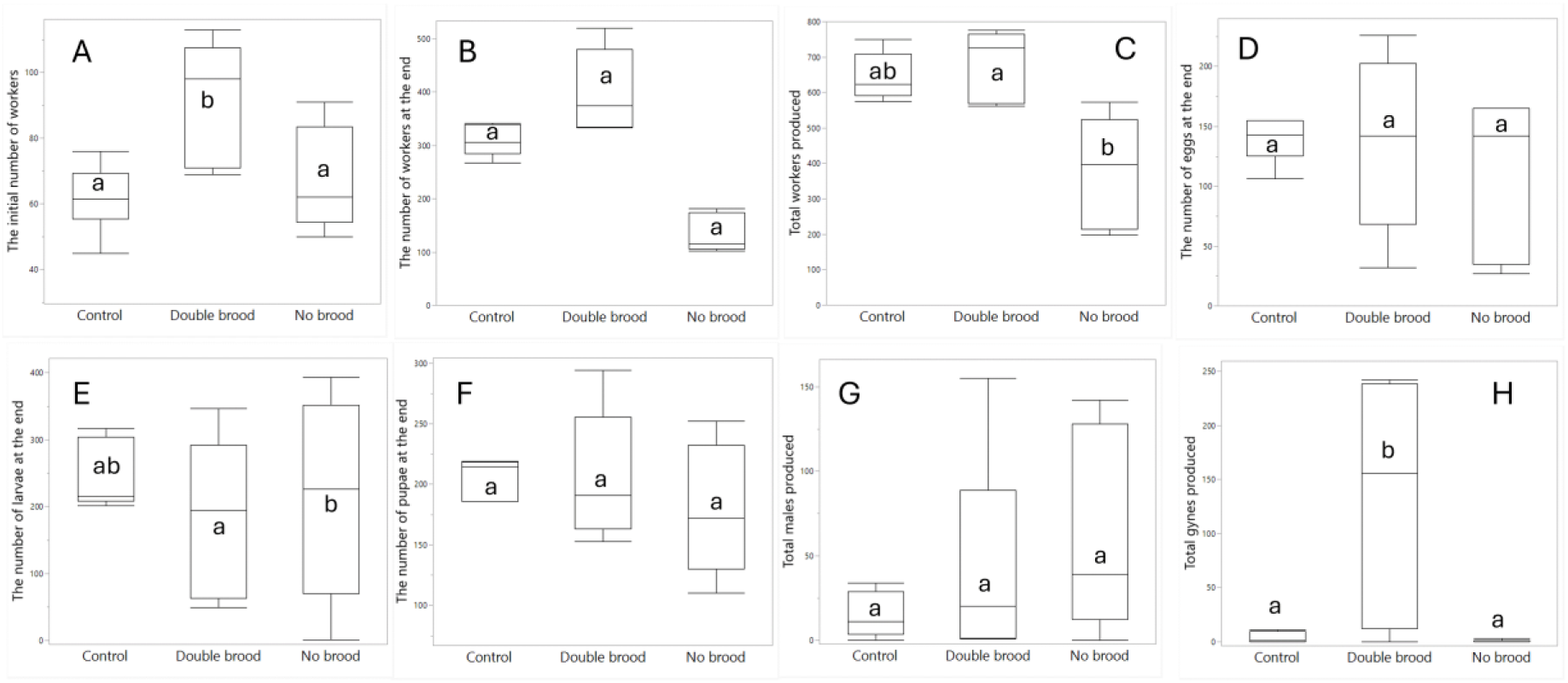
The effect of treatment (no brood, double brood and control) on the initial number of workers (A), the number of workers at the end of the experiment (B), the total number of workers produced (C), the number of eggs (D), larvae (E), and pupae (F) found at the end of the experiment, and the number of males (F) and gynes (H) produced in total. Different letters denote statistical differences at α=0.05.

Treatment had no effect on the number of eggs present at the end of the experiment (F_2,10.4_=0.5, p=0.62, Figure 4D). The number of larvae at the end of the experiment was affected by the treatment (F_2,11_=4.79, 0.03, Figure 4E), with more larvae found in the no brood compared to the double brood (p=0.02). This difference was significantly associated with the total number of workers (F_1,11_=8.47, p=0.01) but not with the initial number of workers (F_1,11_=1.43, p=0.25).

Running the model without the covariates did not converge. The number of pupae at the end of the experiment (Figure 4F) was not affected by the treatment (F_2,10.6_=0.74, p=0.49), but was significantly associated with the total number of workers (F_1,10.6_=6.6. p=0.02).

A treatment-only model excluding covariates showed no effect of treatment on the number of males (F_2,12.5_=2.72, p=0.1, Figure 4G). Adding the total number of workers to the model did not converge, likely due to model complexity combined with the limited sample size. Including only the initial number of workers maintained the non-significance of the model (F_2,12_=0.44, p=0.65) and showed no association with the covariate (F_1,12_=0.001, p=0.97). Including both covariates in the model also failed to converge.

A treatment-only model excluding covariates showed a strong effect of treatment on the number of gynes (F_2,12.2_=8.21, p =0.005, Figure 4H) with double brood colonies producing more gynes than both the no brood (p = 0.007) and control colonies (p = 0.01). Adding the total number of workers as a covariate resulted in a stronger treatment effect (F_2,10.1_=11.1, p=0.002) and no association with the covariate (F_1,10_=2.38, p=0.15). Adding also the initial number of workers to the model did not converge. Including the initial number of workers in the model caused it to fail to converge. Overall, the treatments did not affect the number of eggs, pupae, or males. The number of larvae was affected by treatment but mediated by colony size, whereas the number of gynes was affected by treatment and could not be explained by colony size. We further conducted a survival analysis to compare the timing of gyne production across the three treatments. The Kaplan–Meier survival curves did not differ significantly but approached significance (Log-Rank test: χ² = 4.94, *df* = 2, p = 0.08), suggesting earlier gyne production in the double brood group.

### Aggression by and towards the queen (Figure S1)

There were no differences in the total queen aggression (F_2,12.8_=1.57, p=0.24; Figure S1) or the total worker aggression (F_2,11.9_=1.6, p=0.24). One double brood colony was a clear outlier in the number of the aggressive behaviours performed by and towards the queen on day 4. Thus, we also analysed the data without this colony on day 4 (Figures S1) but this did not change the results.

### Distribution of larva body mass (Figure S2)

The differences between the treatments in gyne production are reflected also in the body mass distribution of larvae that were collected and weighed on the last day of the experiment (day 26). The larvae in double brood colonies showed a bimodal distribution of larva mass, corresponding to larvae that will develop into workers/males and gynes (ACR mode test, p<0.001), while control and no brood colonies showed a unimodal distribution, indicating the production of workers or males (ACR mode test, p>0.114).

### Colony wet mass in colonies restricted to 30/90 workers (Figure 5)

The analysis revealed a significant effect of treatment (F_1,13.9_=11.5, p=0.004), day (F_1,316.7_=382.1, p<0.001) and the interaction between them (F_1,316.7_=20.7, p<0.001). All colonies gained mass over time, reflecting the increase in worker populations and brood. However, colonies with 90 workers gained significantly greater mass over time compared to colonies with 30 workers over the same period.

**Figure 5:**
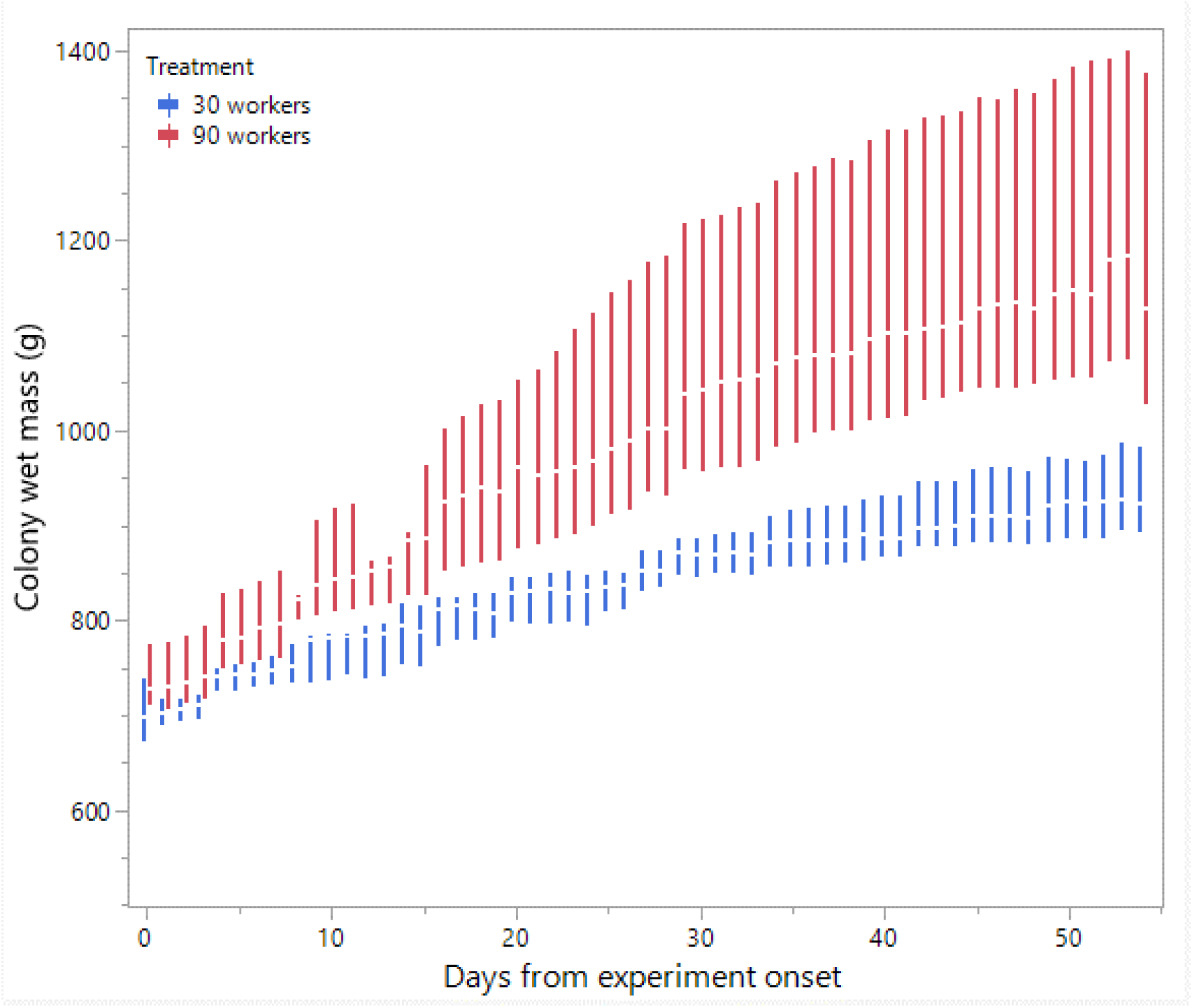
The average colony wet mass throughout the experiment in colonies limited in size to 30 or 90 workers.

### Production of new egg batches in colonies restricted to 30/90 workers (Figure 6)

Neither the treatment (F_1, 13_=0.86, p=0.37) nor the interaction between day and treatment (F_1, 686.8_=0.52, p=0.46) had a significant main effect; however the day alone differed significantly (F_1, 686.8_=71.3, p<0.001), indicating an increase in egg laying over time in both treatments.

**Figure 6:**
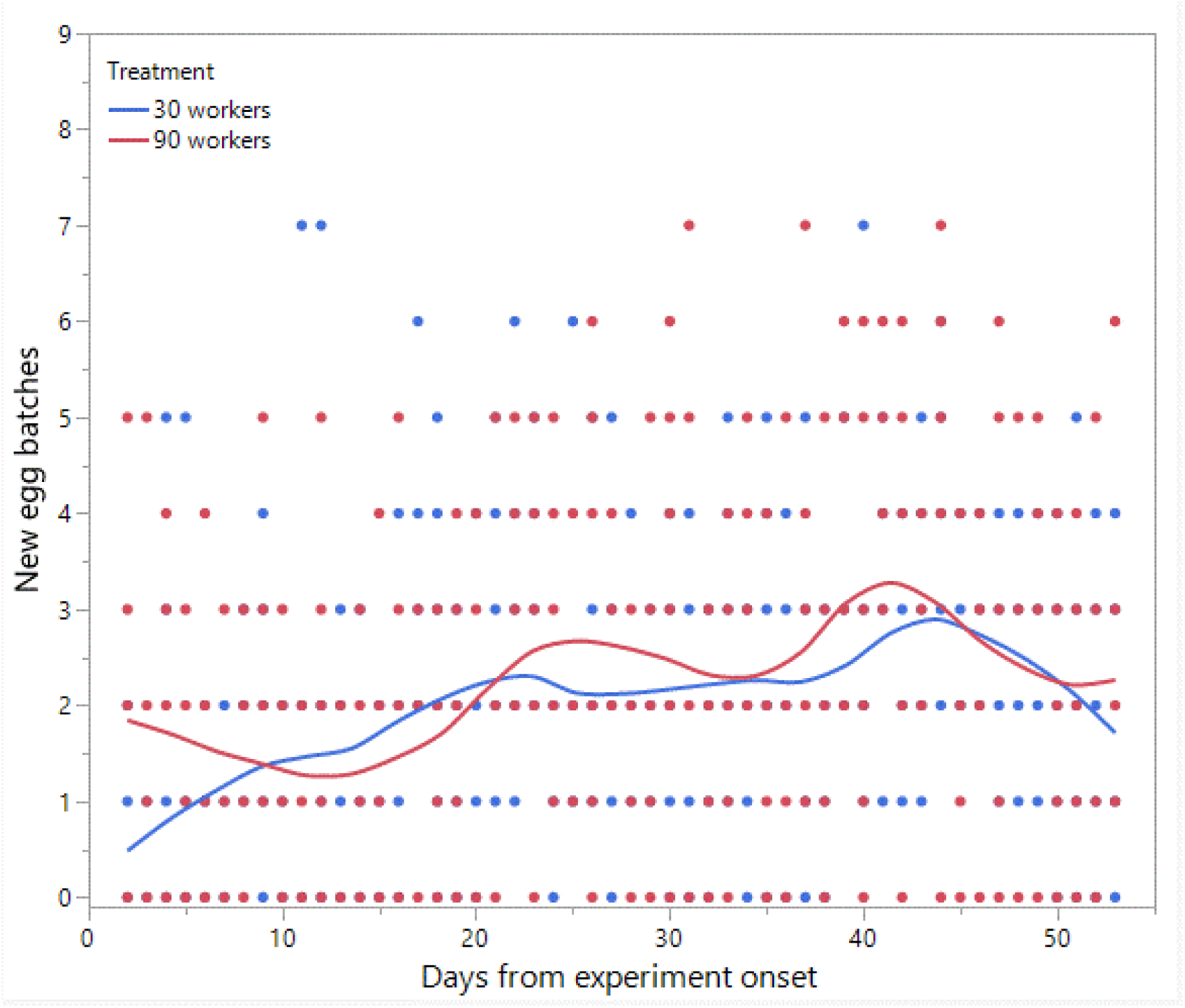
The number of egg batches laid per day throughout the experiment in colonies limited in size to 30 or 90 workers.

### Workers’ ovary activation in colonies restricted to 30/90 workers (Figure 7)

The model showed a significant main effect of treatment (F_1,16.9_= 7.01, p=0.01), but not of day (F_8,639.9_= 1.41, p=0.18) or of the interaction (F_8,639.9_= 1.35, p=0.21), indicating a higher and consistent ovarian activation in the 90-worker treatment.

**Figure 7:**
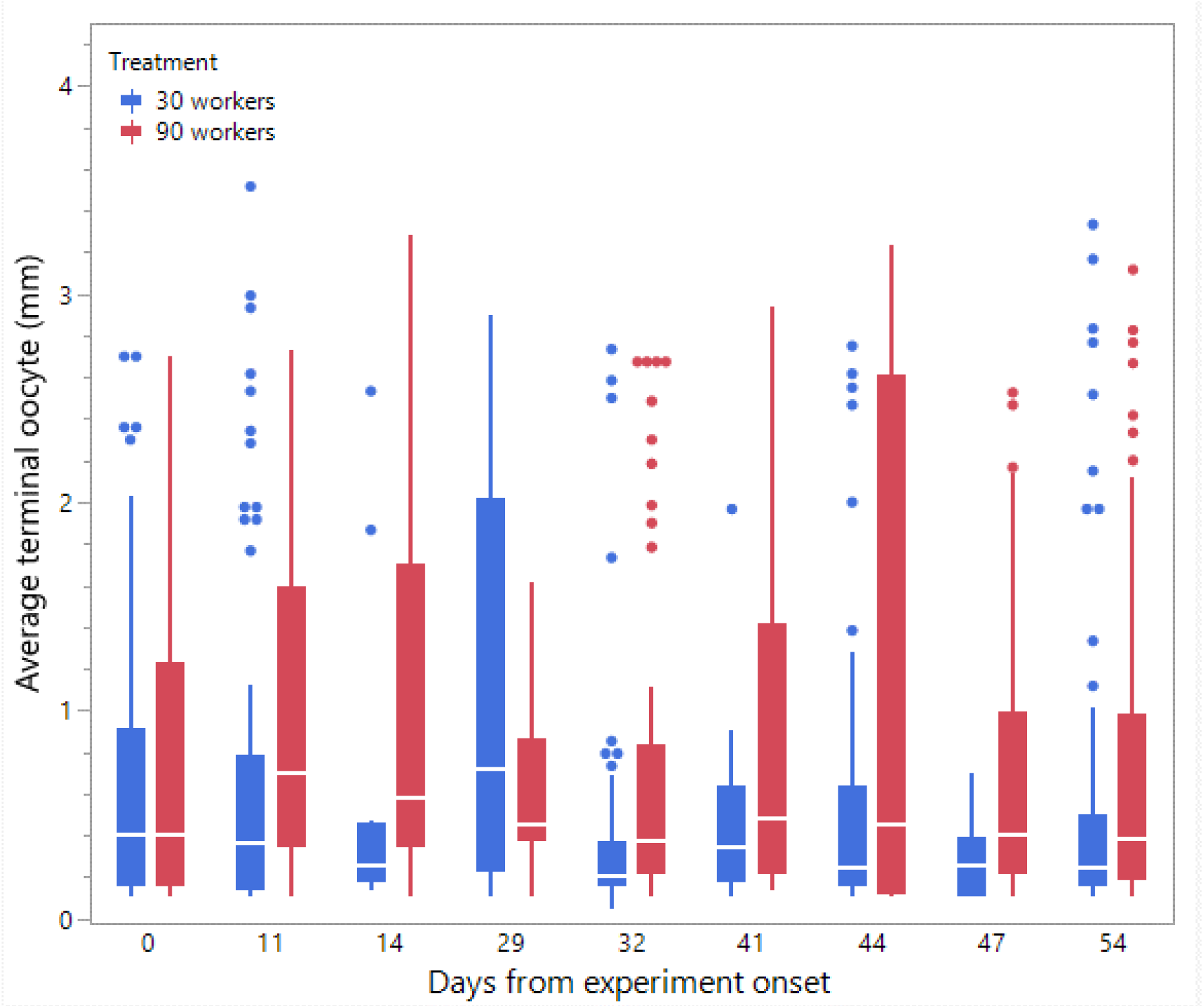
The average terminal oocyte size in workers in colonies restricted to 30 or 90 workers. Oocyte size was measured in nine timepoints throughout the experiment using a subset of workers from each treatment.

### The total number of brood and adults in colonies restricted to 30/90 workers (Figure 8)

The initial mass of brood differed significantly and, despite randomization, was slightly higher in the 90-workers treatment (F_1,13_=6.6, p=0.02, Figure 8A). The number of workers at the end of the experiment was also higher in the 90-worker treatment, in line with our manipulation (F_1,13_=58.3, p<0.001, Figure 8C). While all colonies had more than 30 and 90 workers respectively, the 90- worker colonies had, on average, three times more workers than the 30-worker colonies at the end of the experiment. Neither the initial number of workers prior to the manipulation (F_1,12.9_=1.99, p=0.18, Figure 8B) nor the total number of workers (F_1,13_=0.41, p=0.53, Figure 8D) differed significantly. Therefore, to test the effect of treatment on the number of brood, males, and gynes, we included in the subsequent models both the random effect (colony ID) and the initial brood mass and the end number of workers as covariates.

**Figure 8:**
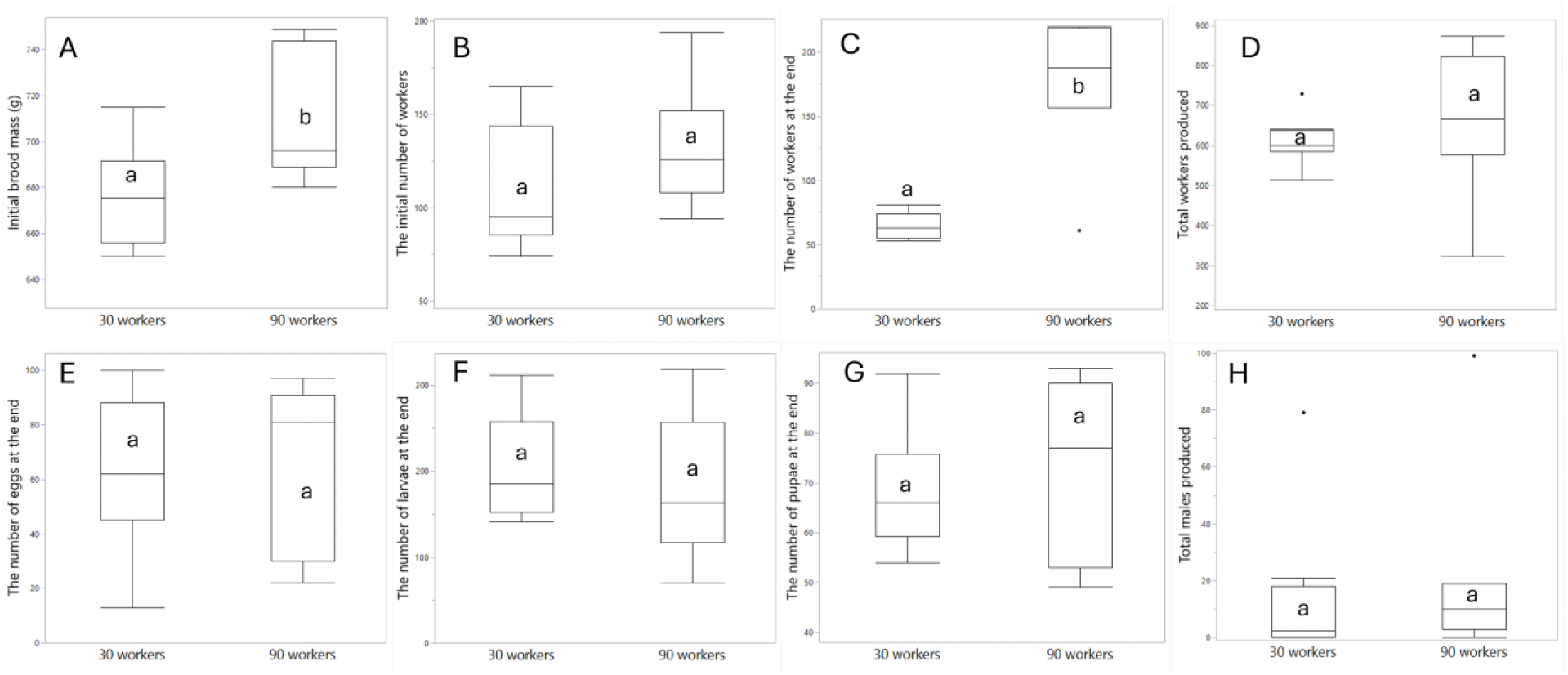
The effect of treatment (30/90 worker colonies) on the initial brood mass (A), The initial number of worker before the manipulation (B), The number of workers found at the end of the experiment (C), the total number of workers produced (D), the number of eggs (E), larvae (F), and pupae (G) found at the end of the experiment, and the number of males (H) produced in total. The total number of gynes produced are not shown since only one colony produced gynes. Different letters denote statistical differences at α=0.05.

The number of eggs present at the end of the experiment was not affected by the treatment (F_1,11_=1.31, p=0.27, Figure 8E) or associated with the initial brood mass (F_1,11_=1.26, p=0.28) or the end number of workers (F_1,11_=3.47, p=0.09). The number of larvae at the end of the experiment was not affected by the treatment (F_1,11_=0.93, p=0.35, Figure 8F) or associated with the initial brood mass (F_1,11_=1.14, p=0.3) or the end number of workers (F_1,11_=1.81, p=0.2).

Similar results were obtained for pupae (Treatment: F_1,11_=0.15, p=0.7; Initial mass: F_1,11_=0.001, p=0.97; End workers: F_1,11_=0.03, p=0.85, Figure 8G). The number of males produced by the colonies did not differ between treatments (F_1,10.4_= 2.77, p=0.12, Figure 8H) and was not associated with any of the covariates (Initial mass: F_1,11_=2.79, p=0.12; End workers: F_1,10.4_=1.14, p=0.3). The number of gynes in the two treatments is not presented since only one colony out of 15 produced any gynes during the experiment. This colony was in the 30-worker treatment and produced 3 gynes in total. All in all, the manipulation of the number of workers did not affect the number of brood or sexuals in the colonies.

### Distribution of larva body mass in colonies restricted to 30/90 workers (Figure S3)

The larvae collected from the colonies at the end of the experiment showed a unimodal distribution indicating the production of workers and/or males only (ACR mode test, p>0.4).

## Discussion

In this study, we demonstrate that manipulating brood quantity in a social colony significantly impacts colony development, gyne production, and worker ovary activation, while showing no significant effect on aggressive behaviour or male production. Differences in gyne numbers between treatments could not be explained by the number of workers, suggesting that brood may function as an extended phenotype of the queen, influencing not only individual reproductive decisions of workers but also colony-level strategies for investing in future queens. Our second experiment, in which colonies were restricted to 30 or 90 workers, unexpectedly resulted in only one colony producing gynes and no significant differences in male production. However, clear differences emerged in worker ovarian activation, with larger colonies showing higher ovary activation of workers. Based on data from both experiments, we suggest that worker ovary activation is a flexible process influenced by colony size, density, or brood amount, whereas sexual production, particularly of gynes, is primarily determined by the amount of brood produced by the colony. Below, we discuss evolutionary explanations for the reliance on sociobiological cues in regulating sexual production, within the framework of optimal resource allocation theory. We also address the evolution of female castes in social insects, along with key caveats and directions for future research.

Optimal resource allocation theories predict that insects secure sufficient resources before switching to sexual production. Evidence across insect species indicates that this transition is influenced not only by external ecological factors but also by internal sociobiological cues (8, 55). These internal cues include queen signalling, queen age, queen presence, and, as demonstrated in this study, brood presence. Such cues are likely more reliable than ecological factors like weather, which may fluctuate, or the number of workers, which reflects past conditions and cannot be reliably extrapolated to predict future ones. Brood may offer a more accurate estimate of ecological resources available for brood rearing, the queen’s fecundity, and the workers’ capacity to support gyne production. Our second experiment demonstrates that colony size alone is not a sufficient cue for gyne production and suggests that a threshold amount of brood is required to trigger its onset. This finding aligns with previous models predicting that bumblebee colonies optimally initiate sexual production approximately one brood development period before colony termination to maximize growth and ensure the successful maturation of sexual offspring (11).

In contrast to gyne production, bumblebee workers appear to rely on multiple cues to initiate male production. These include brood amount and the number of workers as found in this study and also in (31, 56), colony and/or queen age (57), and queen fecundity and health (58). When brood levels are high, workers may delay reproduction to allow for gyne rearing, from which they gain greater indirect fitness benefits compared to males (59). In contrast, the absence of brood suggests that gynes will not be reared, making delayed reproduction unnecessary.

Additionally, the number of workers may signal the approaching end of the season, prompting workers to seize their final opportunity for direct reproduction. This supports the idea that worker reproduction is more flexible and less risky than gyne production, likely because males can be produced by both queen and workers (60) and are less costly to produce .

Previous studies have shown that gyne production and the onset of worker reproduction are highly correlated events (39). Ovarian activation by workers during the competition phase occurs in most colonies, but only some colonies produce gynes. Therefore, these correlations apply only to colonies where both events occur. If the onset of worker reproduction is triggered by various cues - only one of which also triggers gyne production - this could explain both the observed variation among colonies and the correlation between the two events in colonies that exhibit both. Data from other species support this pattern; for example, in *Bombus perplexus*, an increase in worker number is positively correlated with male production, but not necessarily with queen production (61).

A few more points are worth mentioning. First, relatedness could potentially affect the results. All colonies contained non-related workers (as newly emerged workers were distributed daily across all the colonies, including the controls), but only the double brood colonies contained partially unrelated brood, raising the question whether gynes were more likely to develop in colonies with mixed brood. In a previous study we did not find any evidence of the impact of relatedness on brood care (31) and therefore consider it unlikely to have an impact here. We also did not observe any differential behaviour towards the brood in our study in any of the colonies. Whether unrelated brood is more likely to develop into gynes compared to related brood is a question worth investigating. Second, despite our limited number of colonies, we remain highly confident in the effect brood has on the number of sexuals for several reasons: the substantial effect size, the relatively low variation in the number of gynes and males in both the control and no brood colonies compared to the double brood colonies, and our study design which effectively controlled for many confounding factors inherent in the variable nature of a colony.

## Conclusions

Overall, our results show that brood regulates not only egg laying in small groups of workers, as previously demonstrated (31), but also influences colony-level events such as the onset of reproductive competition and the timing of gyne production. We also found that worker ovarian activation is associated with the total number of workers or colony density, even among colonies with similar brood amounts. These findings suggest that brood serves as an indicator of queen fecundity, promoting gyne production when abundant, and possibly triggering the production of cheaper males when absent. In contrast, density alone, though sufficient to induce worker ovary activation, is not a reliable cue for gyne production. Our findings in bumble bees are consistent with results from annual wasps, where partial brood removal increased queen egg laying but did not suppress worker reproduction (62). This may point to a general mechanism in annual social species, in which brood irreversibly influences the level of cooperation between queens and workers and affects sexual production. Based on our second experiment directly testing the effect of density, gyne production likely requires a threshold amount of brood; without it, colonies may instead invest in male production. Indeed, the only colony that produced gynes had 312 larvae at the end of the experiment, while the average in all other colonies was 193.2±19 larvae (mean ± SE). Altogether, our findings highlight the central role of brood in maintaining and shaping social organization in social insects and underscore the need to explore its diverse functions across taxa.

## Declarations

### Availability of data and material

The data used and/or analyzed during the current study are available as supplementary material.

## Acknowledgements

We would like to thank John M. Craig, Andrea Medero and Katie Barie for assisting in the experiments. We also thank the two anonymous reviewers for their thoughtful and constructive comments, which significantly improved the manuscript.

## Author’s contributions

EA, PKFS and EM designed the experiments, PKFS and EM collected the data, CSM performed the behavioral observations, EA analyzed the data, generated the figures and wrote the manuscript. PKFS and EM generated the figures in the supplementary materials. All authors edited the manuscript and gave final approval for publication.

