## Supplementary material for "Gyne production is regulated by the brood in a social bee (*Bombus impatiens*)": Supp material

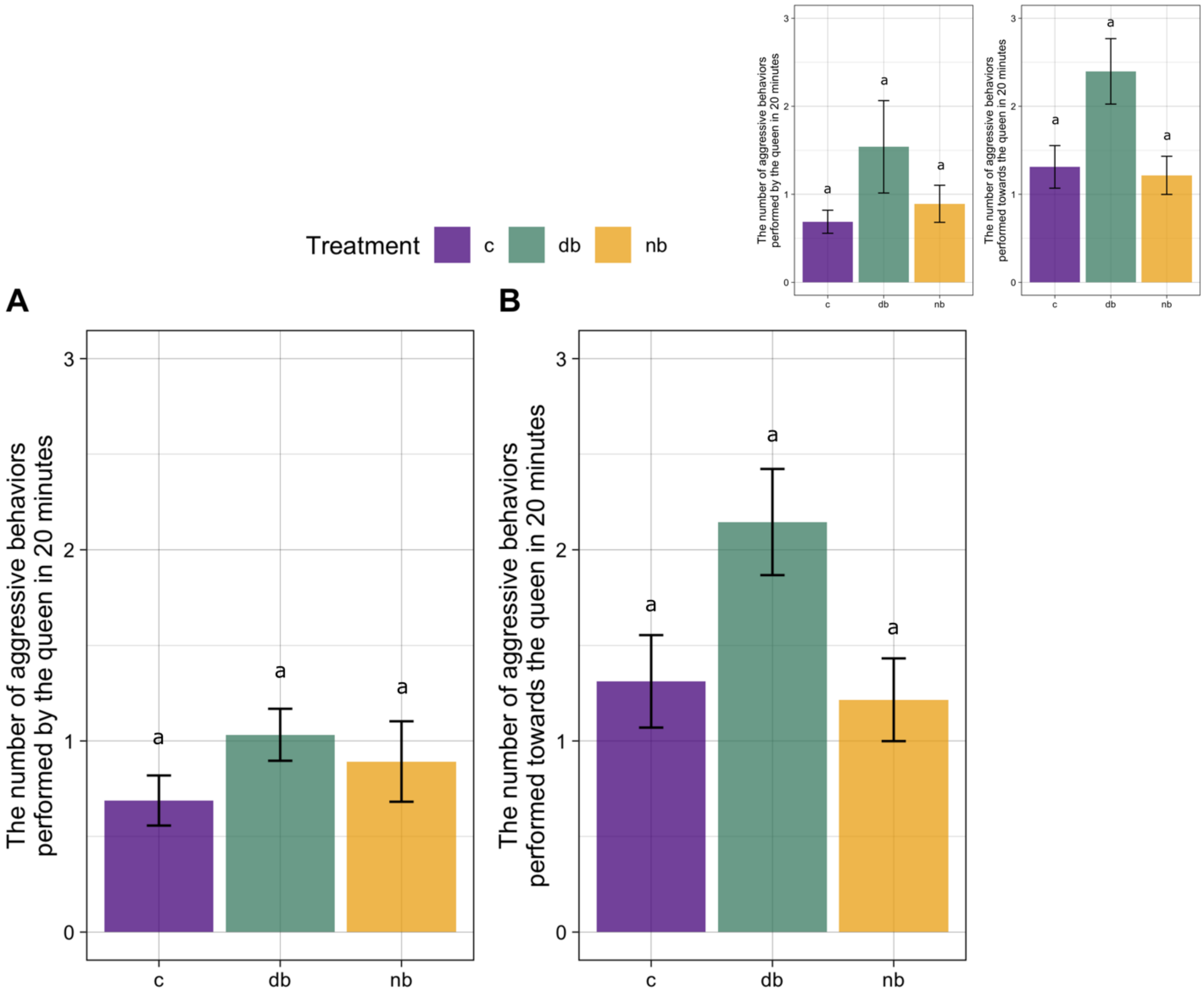


**Figure S1:** The average sum of aggressive behavior events performed towards the queen or by the queen in a 20 min interval across three treatments: no brood, double brood and control. Figures A and B were made using the data without the outliers and the figures on the top right corner using all the data. Data are presented as means ± S.E.M. Different letters denote significant differences at $p < 0.05$.

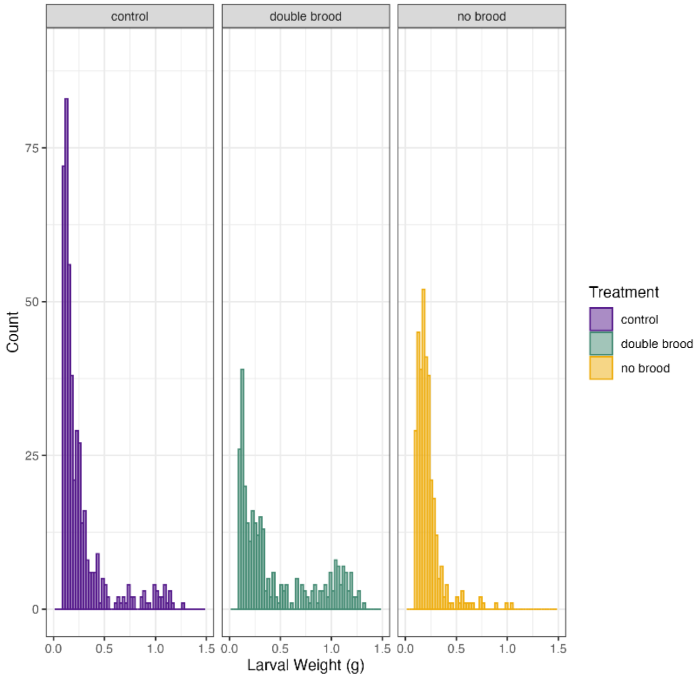


**Figure S2a:** Larval weight distribution across the three treatments (no brood, double brood and control).

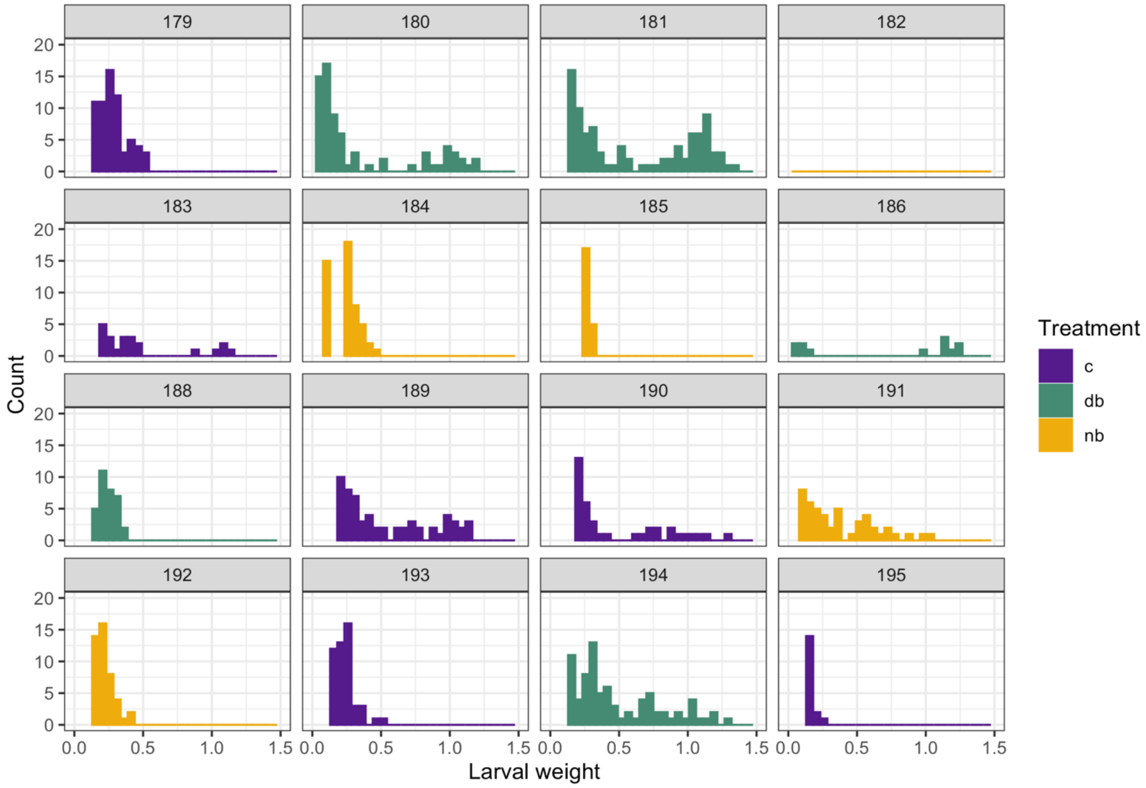


**Figure S2b:** Larval weight distribution by colony across the three treatments: control (c), no brood (nb) and double brood (db).

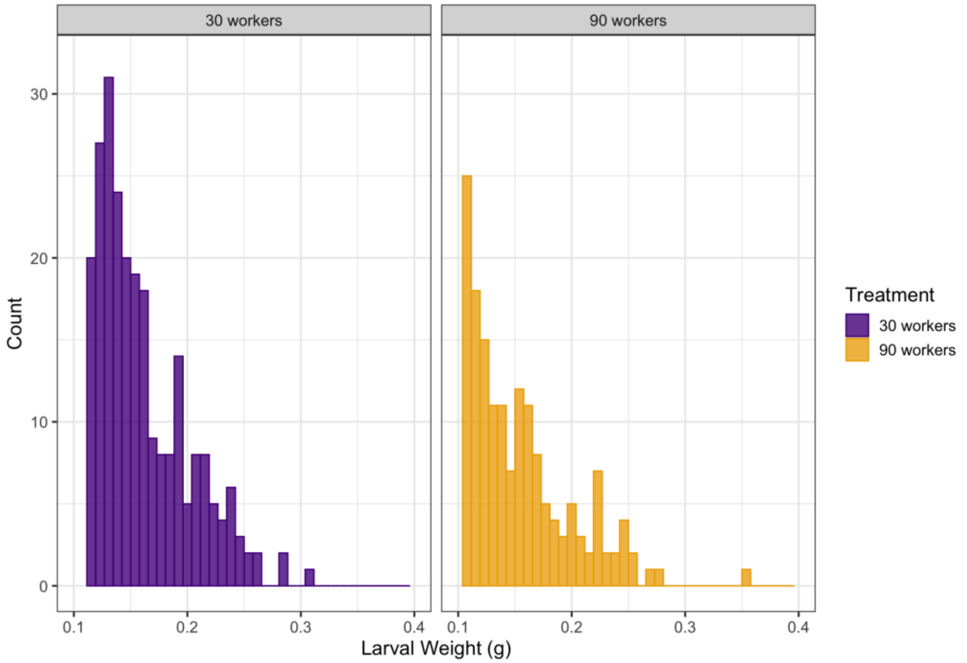

**Figure S3a:** Larval weight distribution across the two treatments (30 and 90 worker colonies).

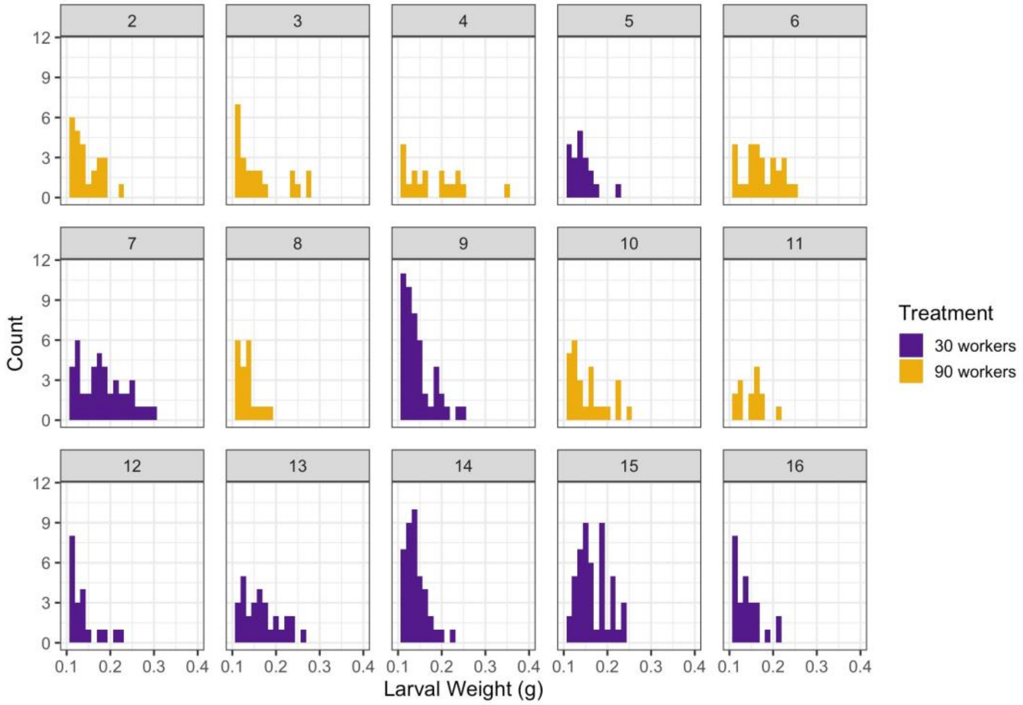

**Figure S3b:** Larval weight distribution by colony across the two treatments: 30 and 90 worker colonies
